# A mean-field model of gamma-frequency oscillations in networks of excitatory and inhibitory neurons

**DOI:** 10.1101/2023.11.20.567709

**Authors:** Farzin Tahvili, Alain Destexhe

**Affiliations:** Institute of Neuroscience (NeuroPSI), Paris-Saclay University, CNRS, 91400 Saclay, France

## Abstract

Gamma oscillations are widely seen in the cerebral cortex in different states of the wake-sleep cycle and are thought to play a role in sensory processing and cognition. Here, we study the emergence of gamma oscillations at two levels, in networks of spiking neurons, and a mean-field model. At the network level, we consider two different mechanisms to generate gamma oscillations and show that they are best seen if one takes into account the synaptic delay between neurons. At the mean-field level, we show that, by introducing delays, the mean-field can also produce gamma oscillations. The mean-field matches the mean activity of excitatory and inhibitory populations of the spiking network, as well as their oscillation frequencies, for both mechanisms. This mean-field model of gamma oscillations should be a useful tool to investigate large-scale interactions through gamma oscillations in the brain.

## Introduction

Information processing in the brain is based on dynamic interplay among different groups of neurons exhibiting various rhythmic patterns. These rhythmic activities of neurons vary significantly depending on different states of behavior and they are closely linked to specific sensory, motor, and cognitive functions. Low-frequency Oscillations (less than 10 Hz) may function as temporal references, as Kayser et al. (2012) suggested. However, faster oscillations within the beta and gamma frequency bands play crucial roles in cognitive processes such as visual attention, feature selection, and binding (Gray et al., 1989; Engel et al., 2001). These oscillatory activities are observed throughout different brain structures across various species and therefore they can be considered as one of the main fingerprints of a typical nervous system.

Among different oscillation bands, the gamma band which is commonly observed during both waking and sleep states, has great importance especially because of its tight relationship to cognitive processes. Generally, oscillations with a frequency higher than 30 Hz are categorized in the gamma band (Buzsáki and Wang 2012). The gamma band can also be divided into sub-bands based on different strategies. For instance, within the hippocampal CA1 region, the application of wavelet analysis revealed the presence of three specific gamma bands, slow gamma (30–50 Hz), mid-frequency gamma (50–90 Hz), and fast gamma (*>* 90 *Hz*) (Tort et al., 2010; Belluscio et al., 2012).

While the collective activity (like the LFP signal) exhibits synchronized oscillations, the individual firing patterns of neurons tend to be irregular and infrequent. This activity regime which is the typical observing state in the cortex in vivo, is known as the Asynchronous Irregular (AI) state and has the following characteristics (Brunel, 2000). This kind of activity places cortical neurons close to their firing threshold rather than resting potential and therefore the response of the system to a stimulus will be much faster than the one at rest. In this regime, neurons are subjected to the bombardment from many inputs so it implies that the excitatory and inhibitory inputs should balance each other and neurons kept around their firing threshold so they fire based on noise fluctuations.

In the presence of stochastic inputs, the asynchronous state may become unstable, leading to the emergence of oscillations (Brunel, 2000; Brunel and Wang, 2003; Geisler et al., 2005). Specifically, it has been shown that these oscillations frequently arise from asynchronous states through Hopf bifurcations when an additional time scale is introduced. This additional time scale goes beyond the one linked to the evolution of the neuron membrane potential and can be attributed to factors such as transmission delay or a finite synaptic timescale (Di Volo and Torcini, 2018). The frequency of these oscillations is typically associated with these external timescales, and since they are in the range of a few milliseconds, this mechanism is commonly associated with fast oscillations.

Previous models demonstrated that gamma oscillations can emerge in a network of inhibitory neurons provided that there is sufficient drive to induce action potentials in the neurons, sufficient strong recurrent connections between them, and at least a time scale apart from the membrane time constant (like the time constant of GABA_*A*_ receptors). In a two-population network of inhibitory interneurons and excitatory pyramidal cells, there can be two possible scenarios for the generation of gamma oscillations, either from within the inhibitory network, or from the loop between inhibitory and excitatory cells. Brunel and Wang (2003) illustrated that in an excitatory-inhibitory network of leaky integrate and fire neurons, the oscillation frequency depends on the synaptic time scales (such as different ionotropic channels’ time constants and synaptic delays) and the relative strength between the excitatory and inhibitory synaptic currents. In particular, they showed increasing the relative strength of excitation versus inhibition typically decreases the frequency of oscillation.

Special characteristics of gamma oscillations are as follows (Buzsáki and Wang, 2012). To begin with, gamma oscillations are of brief duration and arise from the interplay between excitatory and inhibitory processes. Additionally, various sub-bands of gamma oscillations can coexist or manifest independently. Furthermore, the genesis of these rhythms is closely linked to inhibitory processes. Moreover, their occurrence typically aligns with the irregular firing of individual neurons, and the network frequency of gamma oscillations varies widely based on the underlying mechanism. Lastly, the amplitude of gamma oscillation is influenced by slower rhythms.

Gamma oscillations have been found in several areas of the brain such as neocortex (Sirota et al., 2008), entorhinal cortex (Chrobak and Buzsaki, 1998), amygdala (Popescu et al., 2009), hippocampus (Mann et al., 2005), striatum (Berke et al., 2004), and thalamus (Pinault and Deschênes, 1992). Common properties of these brain regions are the presence of inhibitory interneurons and their actions through GABA_*A*_ receptors. The consistent patterns of gamma frequency oscillations across various brain regions and species offer insights into the necessary mechanisms that support them.

In this study, we work with conductance-based sparse spiking networks that consist of two types of neurons, pyramidal regular-spiking (FS) neurons and fast-spiking (FS) interneurons, and investigate the genesis of gamma rhythms in these RS-FS networks. We offer the corresponding mean-field model in parallel and illustrate that it perfectly agrees with the spiking networks. In particular, we investigate the role of synaptic delay with specific attention due to its proved important role (Brunel and Wang, 2003).

We start by investigating the gamma oscillation emergence during the AI state and provide the mean-field model which is in very good agreement with spiking networks. We next investigate two mechanisms that result in gamma oscillations appearing, synaptic delay and excitation-inhibition balance break. In all cases, we compare spiking network models with their corresponding mean-field, and discuss in which experimental conditions they can be realistic. Finally, we discuss experimentally observed characteristics of gamma oscillations that were mentioned before, and how they can be observed in both types of models.

## Materials and Methods

### Spiking Network Model

We use the AdEx model for the dynamics of our neurons. Each neuron is described by its membrane potential *V* and the adaptation current *w*, which the dynamics are as follows.

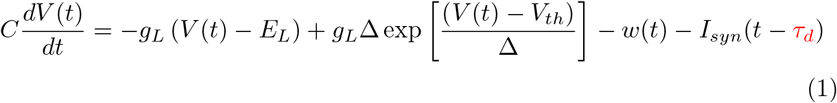

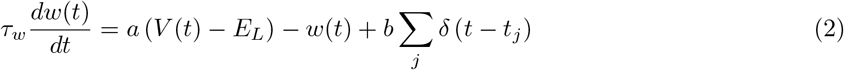

Where *C* is the neuron membrane capacitance, *g*_*L*_ is the leakage conductance, *E*_*L*_ is the leaky membrane potential, *V*_*th*_ is the effective threshold, and Δ is the threshold slope factor. *τ*_*w*_ is the adaptation time scale, *a* indicates the sub-threshold adaptation factor, and *b* is the spike-triggered adaptation factor. The synaptic current, *I*_*syn*_(t), is the sum of all currents that are injected into the neuron due to the spiking activity of other cells. *τ*_*d*_ is a synaptic time delay.

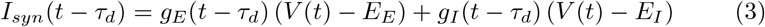

*E*_*E*_ and *E*_*I*_ are the reversal potential of excitatory and inhibitory synapses, respectively. We take the synapses as conductance-based which means every time a pre-synaptic neuron spikes at time *t*_*k*_, the excitatory *g*_*E*_ or the inhibitory *g*_*I*_ synaptic conductance increases by a discrete amount *Q*_*E*_ or *Q*_*I*_ (excitatory or inhibitory synaptic quantal), depending on the nature of the pre-synaptic neuron. We consider the dynamics of the conductances as follows.

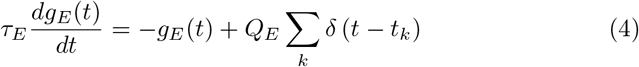

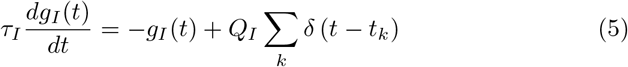

*τ*_*E*_ and *τ*_*I*_ are the time scales of the excitatory and inhibitory synapses, respectively. The Σ_*k*_ runs over all the presynaptic excitatory or inhibitory neurons spike times. During the simulations, the equations characterizing the membrane potential *V* and *w* are integrated until a spike is generated. In practice, the spiking time is defined as the moment in which *V* reaches a certain threshold (*V*_cut_). Then the membrane potential is reset to *V*_rest_, which is kept constant until the end of the refractory period *T*_ref_. After the refractory period, the equations start being integrated again. The adaptation current increases by an amount *b* every time that the neuron emits a spike at times *t*_*j*_. For the specific values of the parameters, see Table 1 and Table 2 (the parameter values have been extracted from Destexhe, 2009 and Destexhe et al., 1998).

**Table 1.**
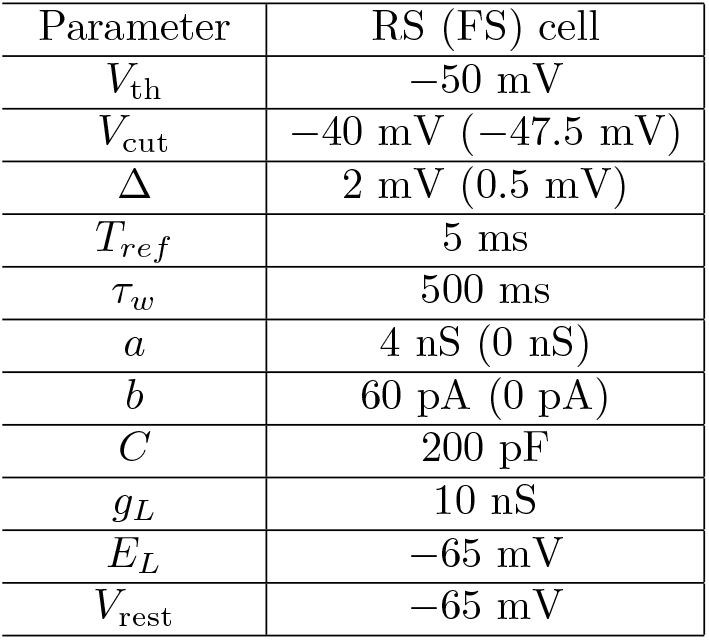
Neuron parameters.

| Parameter | RS (FS) cell |
| --- | --- |
| $V_{th}$ | −50 mV |
| $V_{cut}$ | −40 mV (−47.5 mV) |
| $\Delta$ | 2 mV (0.5 mV) |
| $T_{ref}$ | 5 ms |
| $\tau_w$ | 500 ms |
| $a$ | 4 nS (0 nS) |
| $b$ | 60 pA (0 pA) |
| $C$ | 200 pF |
| $g_L$ | 10 nS |
| $E_L$ | −65 mV |
| $V_{rest}$ | −65 mV |

**Table 2.** Synapse parameters.

| Parameter | Type 1 (Type 2) |
| --- | --- |
| $\tau_E$ | 5 ms |
| $\tau_I$ | 5 ms |
| $E_E$ | 0 mV |
| $E_I$ | −80 mV |
| $Q_E$ | 1.25 nS (1 nS) |
| $Q_I$ | 5 nS |
| $\tau_d$ | *Free* |
| $\epsilon$ | 0.05 (0.03) |

We consider two types of synapses between our neurons, excitatory and inhibitory synapses. For determining the time scale of the excitatory synapse, we consider that such a synapse is composed of 10% NMDA receptors and the rest (90%) AMPA receptors and this leads to a net time scale equal 5 ms. For the inhibitory synapses which we suppose only include GABA_*A*_ receptors, the time scale is considered equal to 5 ms (Destexhe et al., 1998). Table 2 shows synaptic parameters including synaptic conductance quanta *Q*_*E*_ and *Q*_*I*_, synaptic time scales *τ*_*E*_ and *τ*_*I*_, and synaptic delay *τ*_*d*_ and parameter *ϵ*.

We consider synaptic time delays (*τ*_*d*_) in the synaptic interactions between neurons. In other words, we assume that when a pre-synaptic neuron fires an action potential, it takes some time *τ*_*d*_ to be received by the post-synaptic cell. In the spiking network, we can simply suppose that the delay for all kinds of connections (*exc→exc, inh→exc, exc →inh*, and *inh→inh*) be the same. However, in the mean-field model, we need an asymmetry in delays to generate the oscillations and due to this reason, we consider an asymmetry of time delays (both in the spiking network and the mean-field), using an asymmetry parameter *ϵ <<* 1, as follows: The delays for *exc → exc* and *inh → inh* connections are set to (1 + *ϵ*)*τ*_*d*_ while the delays for the other two connections (*exc → inh* and *inh → exc*) are set to (1*−ϵ*)*τ*_*d*_. For more explanations about this asymmetry, we refer the reader to Appendix A. The value of *ϵ* which are used in simulations is 0.05 and 0.03 for Type 1 and Type 2 networks, respectively (see Results).

Our networks are composed of *N* = 10000 neurons, 80% regular spiking, and 20% of inhibitory fast-spiking cells. All neurons are connected randomly with the probability connection, *p*, which we take it equal to 0.05 . In addition to recurrent connections, neurons also receive an external drive. The drive is implemented as 8000 independent and identically distributed Poissonian spike trains with the spiking frequency *μ* (The typical value of *μ* in our simulations is 4 *Hz*), being sent to the network with the probability connection *p*.

Throughout the study, we determine the population activity of the spiking network as follows: We employ the population activity definition,

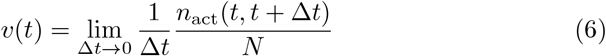

which allows us to calculate the population activity for both the RS and FS populations after simulating the spiking network. In this formula, N represents the number of neurons within the specific population, such as FS and RS populations. The term *n*_*act*_(*t, t* + Δ*t*) signifies the cumulative spike count across all neurons in the group within the time window (*t, t* +Δ*t*). Throughout our study, we consistently set Δ*t* to 0.1 *ms* for calculating spiking network population activities. After calculating the population activities, we perform Fourier analysis on them and plot the spectrum of the signals. We choose the frequency which has the maximum power as the frequency of oscillation.

### Mean-Field Model

The mean-field model consists of a model introduced previously (Di Volo et al. 2019), which consists of two populations of RS and FS neurons with adaptation. This model was based on a Master Equation formalism to derive mean-fields of spiking neurons (El Boustani and Destexhe, 2009), and which was extended to model networks of RS and FS cells described by the AdEx model. The version of Di Volo et al. (2019) included adaptation and closely matched the spontaneous activity as well as the response of the network to external inputs. The main modification to this model is that we introduced time delays to match the synaptic delay in the spiking network. Our mean-field dynamics are as follows:

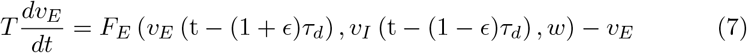

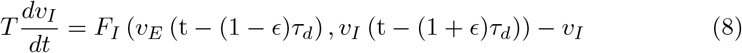

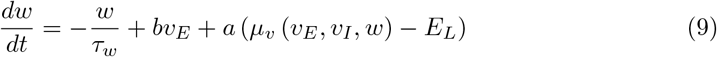

*v*_*E*_ and *v*_*I*_ are the mean population activities of RS and FS populations, respectively. *F*_*E*_ and *F*_*I*_ are RS and FS transfer functions, respectively. *μ*_*v*_ is the RS average population voltage which is a function of *v*_*E*_, *v*_*I*_, and *w* (the formulation of transfer functions and *μ*_*v*_ are provided in Appendix B). *τ*_*d*_ is the time delay that mimics the role of synaptic delay in the spiking network and *ϵ* is the parameter explained in the spiking network part. *w* is the adaptation factor and a and b are adaptation parameters same as in the spiking network. *E*_*L*_ is the leak voltage which equals that of the spiking network. The equation for the FS population has no term *w* since the adaptation of FS neurons is considered to be zero. T is the time scale of the mean-field dynamics. On the principles of the Master Equation formalism (El Boustani and Destexhe, 2009), T must be in the range in which the system acts as a Markovian system. However, if we put it high, the mean-field cannot capture gamma oscillations. Throughout the study, we consider T = 5 ms and so we ensure that we can capture all the frequencies below 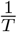 which is 200 Hz.

We use the Ornstein-Uhlenbeck process as the noise in our mean-field model which is the only nontrivial process that satisfies Gaussian, Markovian, and temporally homogeneous properties. The noise is defined by the following equation *dX*(*t*) = *−X*(*t*)*θdt*+*σdW* (*t*), where *σ* and *θ* are parameters and *W* (*t*) is the standard Brownian motion. We take the timescale of the noise equal to 5 *ms*, resulting in choosing *θ* = 200 *Hz*. This amount of timescale is the same as in Di Volo et al. (2019), where they model Up and Down states. By doing this, on the one hand, we ensure that all frequencies below 200 *Hz* exist in the noise with comparable power, and on the other hand, it is completely compatible with the configuration of the mean-field which explains Up-Down states in Di Volo et al. (2019).

### Simulation

Spiking networks are simulated under the Brian 2 Python package and the mean-fields are simulated under Python. All equations are numerically integrated using Euler’s method with *dt* = 0.1*ms* as the integration time step (smaller time steps also lead to similar results).

## Results

In this section, we investigate two types of networks where gamma oscillations can appear, which we call Type 1 and Type 2. From the perspective of parameters, the only major difference between them is that in Type 1 the external drive on both populations is equal, while in Type 2 the external drive on the RS population is higher than that of the FS population. We model them in the spiking network and show that the mean-field matches well with the spiking network in both cases.

### Type 1 networks

In Type 1 networks, we consider the situation in which the external drive is the same for both populations and we take it equal to 4 *Hz*. We illustrate that in this case, the synaptic delay plays a major role in the generation of gamma oscillations.

Figure 1 shows the dynamics of the spiking network with no delay, the meanfield, and their frequency analysis. Only very fast oscillations can be seen from the raster plot or RS and FS population activities. This is also confirmed by the frequency analysis of the network activity. The corresponding mean-field is in excellent agreement with the firing rates of the spiking network.

**Figure 1.**
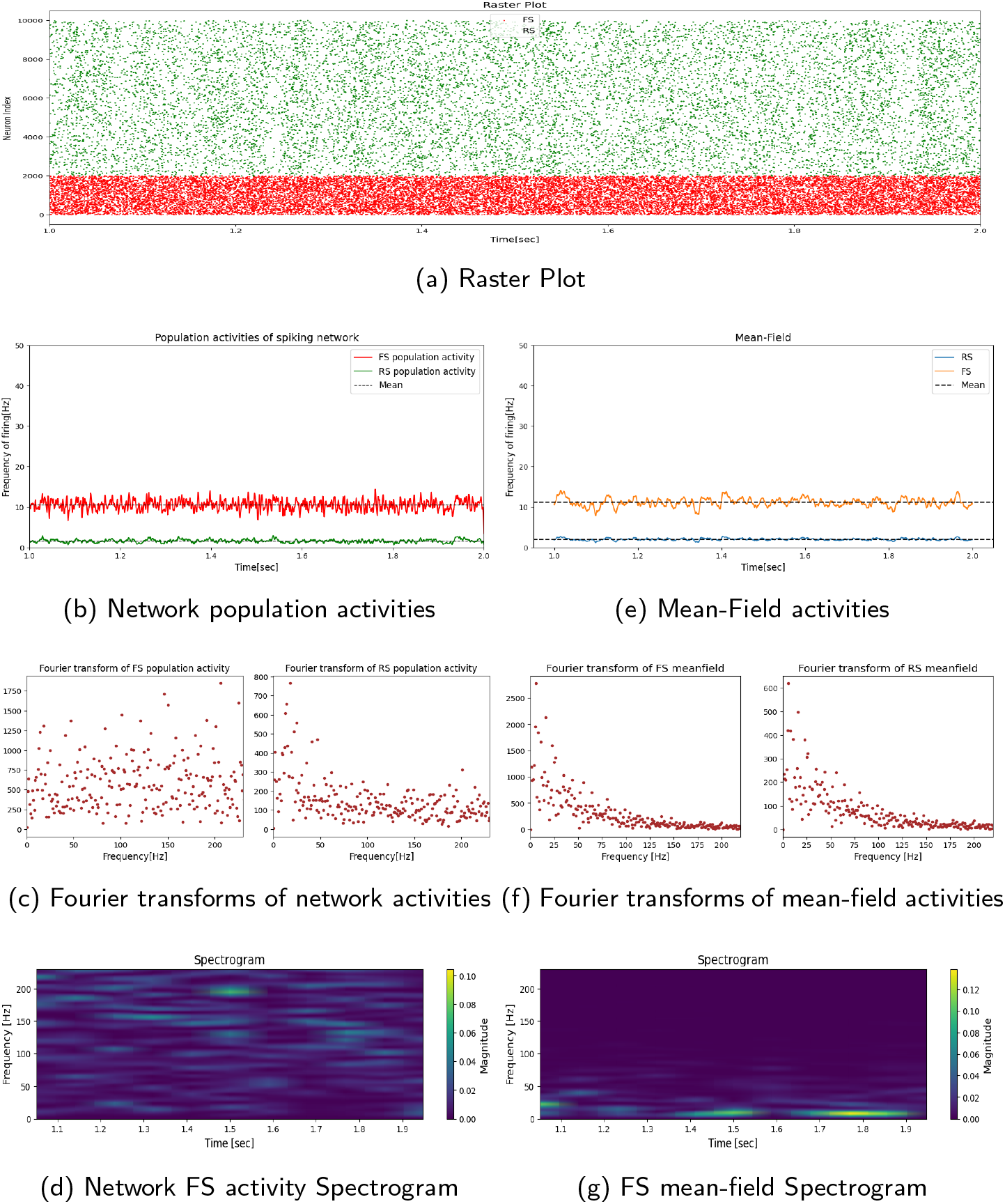
Type 1 spiking network and its corresponding mean-field without delay. Figure (a) shows the simulated raster plot of the spiking network over a one-second interval. Figure (b) shows the spiking network’s RS and FS population activities. Figure (c) shows the Fourier transforms of activities in figure (b), and figure (d) is the Spectrogram of the FS activity of the figure b. Figures (e), (f), and (g) are the same as figures (b), (c), and (d) respectively, but for the mean-field. By comparing panels b and e shows a good match between the spiking network and the mean-field in terms of mean activities. This can also be seen from the Fourier transform plots of both the network and the mean-field, (and also Spectrograms), where no obvious dominant peak is visible (apart from noise) thus the activities are not considered as oscillatory.

The situation is different when one considers a synaptic delay in the system. The spiking network in Fig. 2 clearly shows a prominent oscillation and the oscillation frequency is around 100Hz in both populations. The corresponding mean-field for a delay of 2 ms is shown in the same figure, Fig. 2, and it matches very well the spiking network in terms of both the frequency of oscillation and the mean activities. Also the power spectrum and Fourier transforms of the mean-field to those of the network, shows that the power of oscillation in the FS population is higher than that of RS, similar to the spiking network.

**Figure 2.**
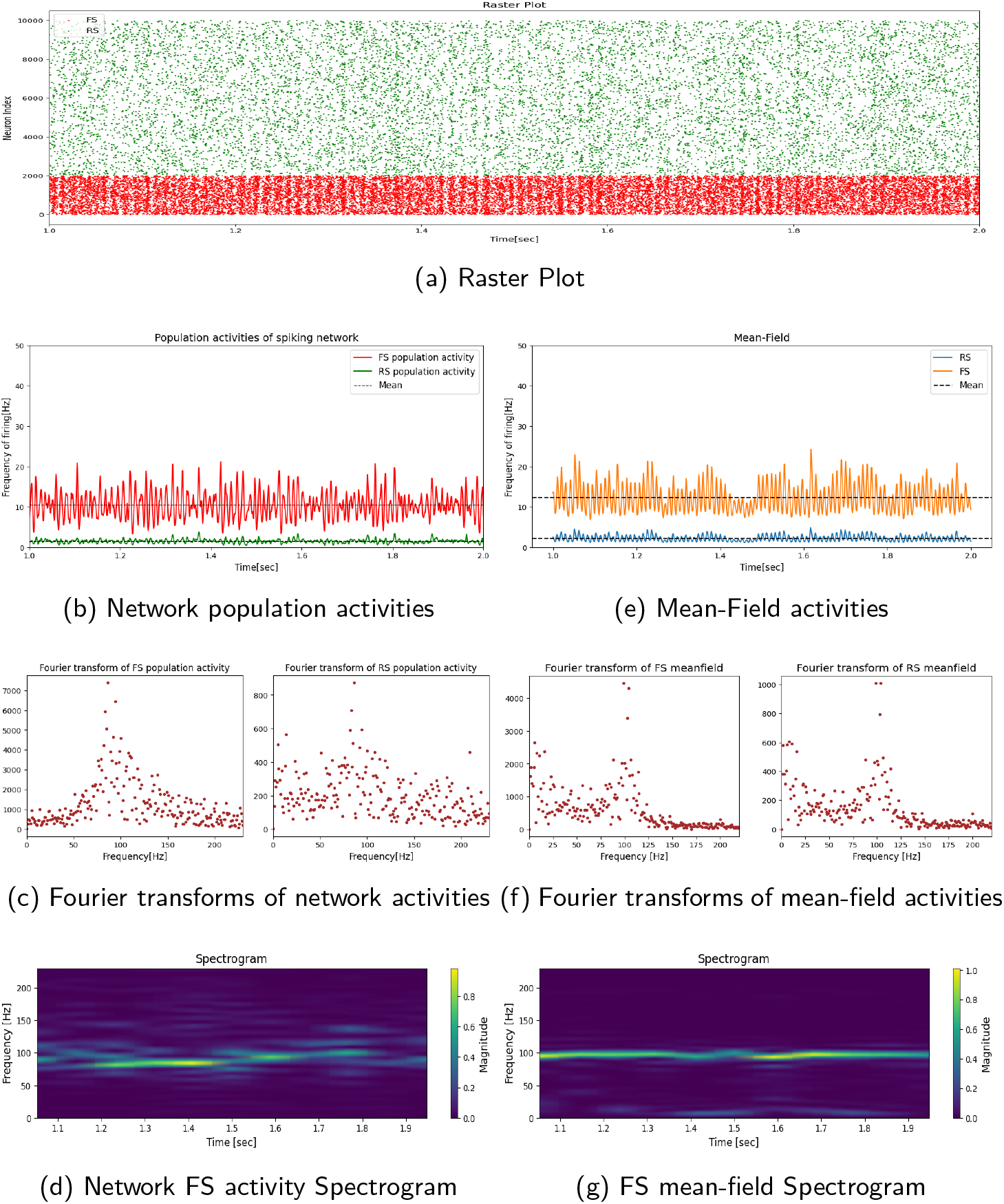
Type 1 spiking network and its corresponding mean-field with 2 ms delay. Same arrangement of panels as for Fig. 1. The system has a prominent oscillation of around 100 Hz which is nearly the same in the spiking network and the mean-field as seen through Spectrograms and Fourier transform curves. Furthermore, the power of oscillations in the network is in agreement with that of mean-field. Additionally, the power of oscillation in the FS population is higher than that of the RS population both in the network and mean-field. Overall, The mean-field model is in good agreement with the spiking network regarding both population mean values and the oscillation characteristics.

The same observations hold for a delay of 3 *ms* (Fig. 3). In this case, the oscillation frequency is around 75 Hz, and the mean-field matches well with the spiking network in terms of both oscillation frequency and the mean activities of RS and FS cell populations. Moreover, one can see that when we increase the delay, the synchrony increases, and the magnitudes of oscillations become larger (compare Fig. 3 with Fig. 2).

**Figure 3.**
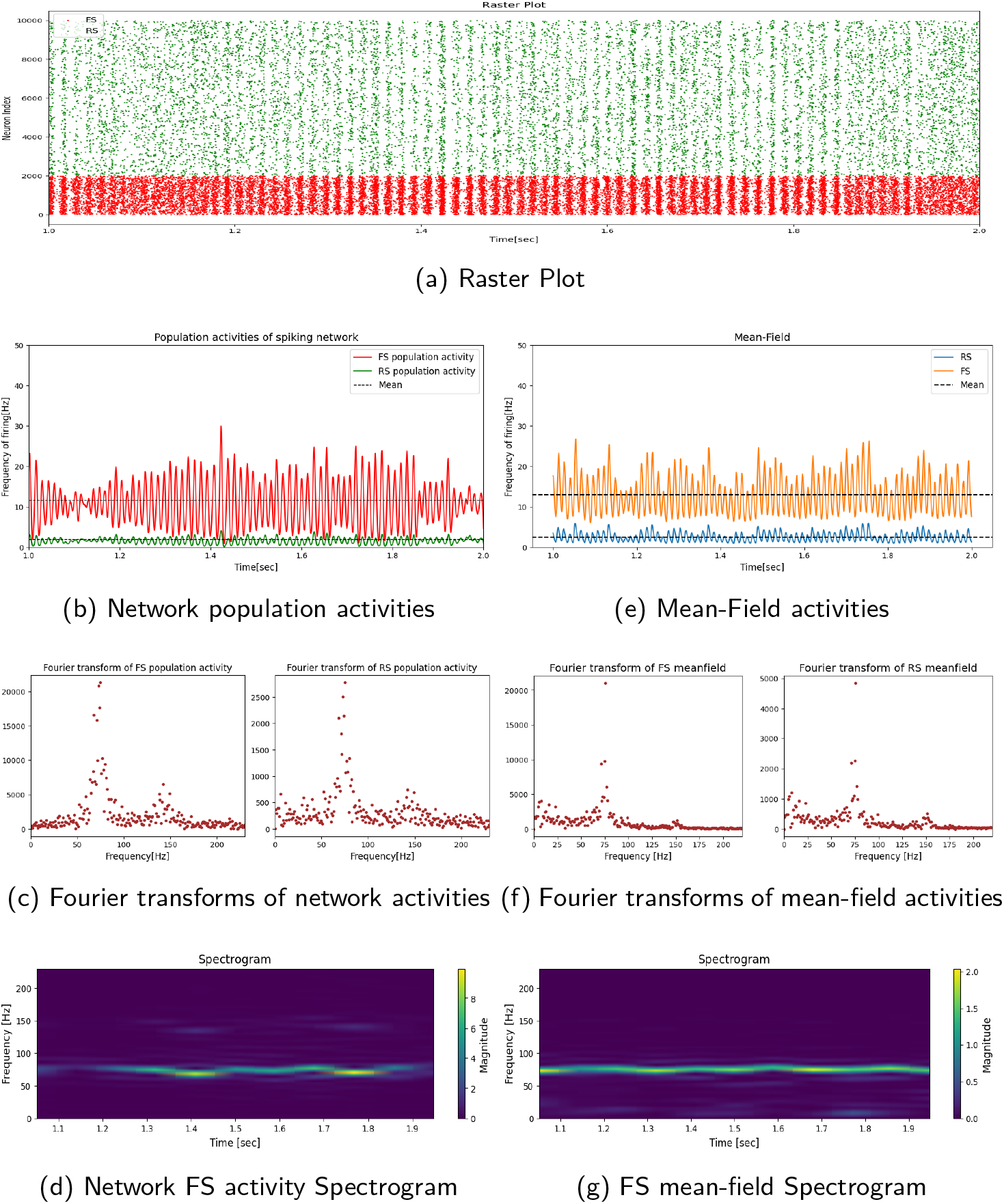
Type 1 spiking network and its corresponding mean-field with 3 ms delay. Same arrangement of panels as for Fig. 1. The system has a prominent oscillation of around 75 Hz which is nearly the same in the spiking network and the mean-field as seen through Spectrograms and Fourier transform curves. Furthermore, the power of oscillations in the network is in agreement with that of mean-field. Additionally, the power of oscillation in the FS population is higher than that of the RS population both in the network and mean-field. Overall, The mean-field model is in good agreement with the spiking network regarding both population mean values and the oscillation characteristics.

So, for Type 1 networks, we conclude the following. First, as one increases the synaptic delay, the network synchrony, the magnitude of oscillations, and the ratio of gamma power are amplified (compare the power of oscillations of Fig. 1, Fig. 2 and Fig. 3 shown besides Spectrograms or the y-axis of Fourier plots). Secondly, the Power of gamma in the FS population is much higher than that of the RS population (compare the Fourier spectra of different cell type activities). Figure 4 shows a synthesis of the network simulations and the corresponding mean-field. One can see that there is an excellent agreement between the spiking network and its mean-field in terms of the frequency of oscillation. Moreover, it is obvious from the figure that the frequency of oscillation is a decreasing function of synaptic time delay.

**Figure 4.**
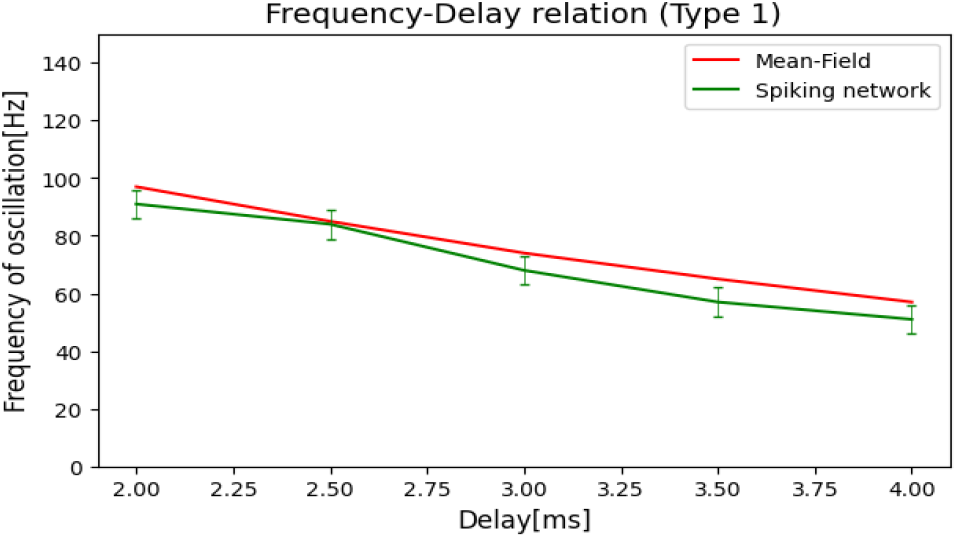
Frequency of oscillation as a function of synaptic delay in Type 1 networks. The mean-field frequency matches well with that of the spiking network. Moreover, the oscillation frequency as a function of synaptic time delay is a decreasing function.

### Type 2 networks

In this case, we consider the situation in which we can apply a lower external drive on the FS population compared to that of the RS population. This case could correspond to drugs that would decrease the synaptic input to FS cells, or to brain areas where the external input is weaker on FS cells. In such cases, we artificially break the excitation-inhibition balance. To be specific, we apply external drives of 2 *Hz* on the FS and 4 *Hz* on the RS populations (in Type 1 it was 4 *Hz* for both populations). Other parameter values that differ from those of Type 1 are *Q*_*e*_ which we take it 1 *nS* instead of 1.25 *nS* and also here in Type 2 we take *ϵ* = 0.03.

Indeed, when simulating Type 2 RS-FS spiking networks, we observe the genesis of gamma oscillations, as shown in Figs. 5, 6, 7, and 8. In this case, the power of oscillation is comparable for RS and FS cells and similar to Type 1 networks, there is a good match between the spiking network and the mean-field in terms of both frequency of oscillation and mean activity values, as one can see by comparing spiking networks and the mean-field in Fig. 5, 6, 7, and 8. Thus, there is a good general agreement between the mean-field and the spiking network for Type 2 as well. The frequency-delay relation is further shown in Fig. 9, where the gamma oscillation frequency predicted by the mean-field closely matches the frequency of the Type 2 spiking network. Another important point that one should pay attention to, is that in Type 2 the presence of delay is not necessary for the generation of gamma oscillation, as one can see that the system with no delay can also generate some oscillations in the gamma range (see Fourier transforms in Fig. 5). However, adding the time delay to the system causes the oscillations more prominent with a much higher power which is easily distinguishable from noise (compare Fig. 5 with Figs. 6, 7, and 8).

**Figure 5.**
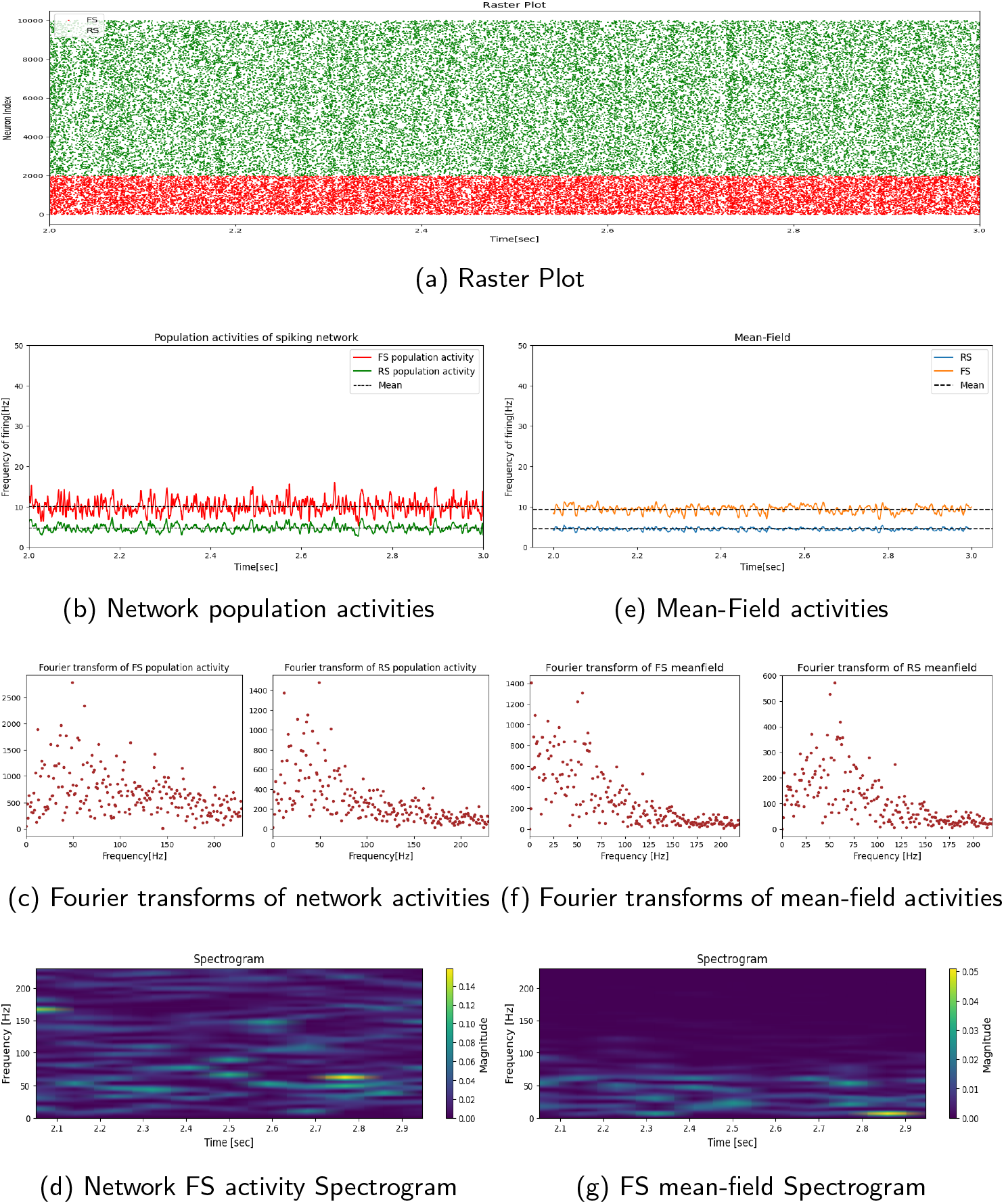
Type 2 spiking network and its corresponding mean-field without delay. The external inputs on excitatory and inhibitory populations are 4 Hz and 2 Hz, respectively. Same arrangement of panels as for Fig. 1. Comparing figures b and e shows a good match between the spiking network and the mean-field in terms of mean activities. In the Fourier transform plots of both the network and the mean-field, one can see a peak around 50 Hz but their powers are weak compared to the same cases with delay (compare this figure with figures 6, 7, and 8). Overall, the mean-field and network are in good agreement.

**Figure 6.**
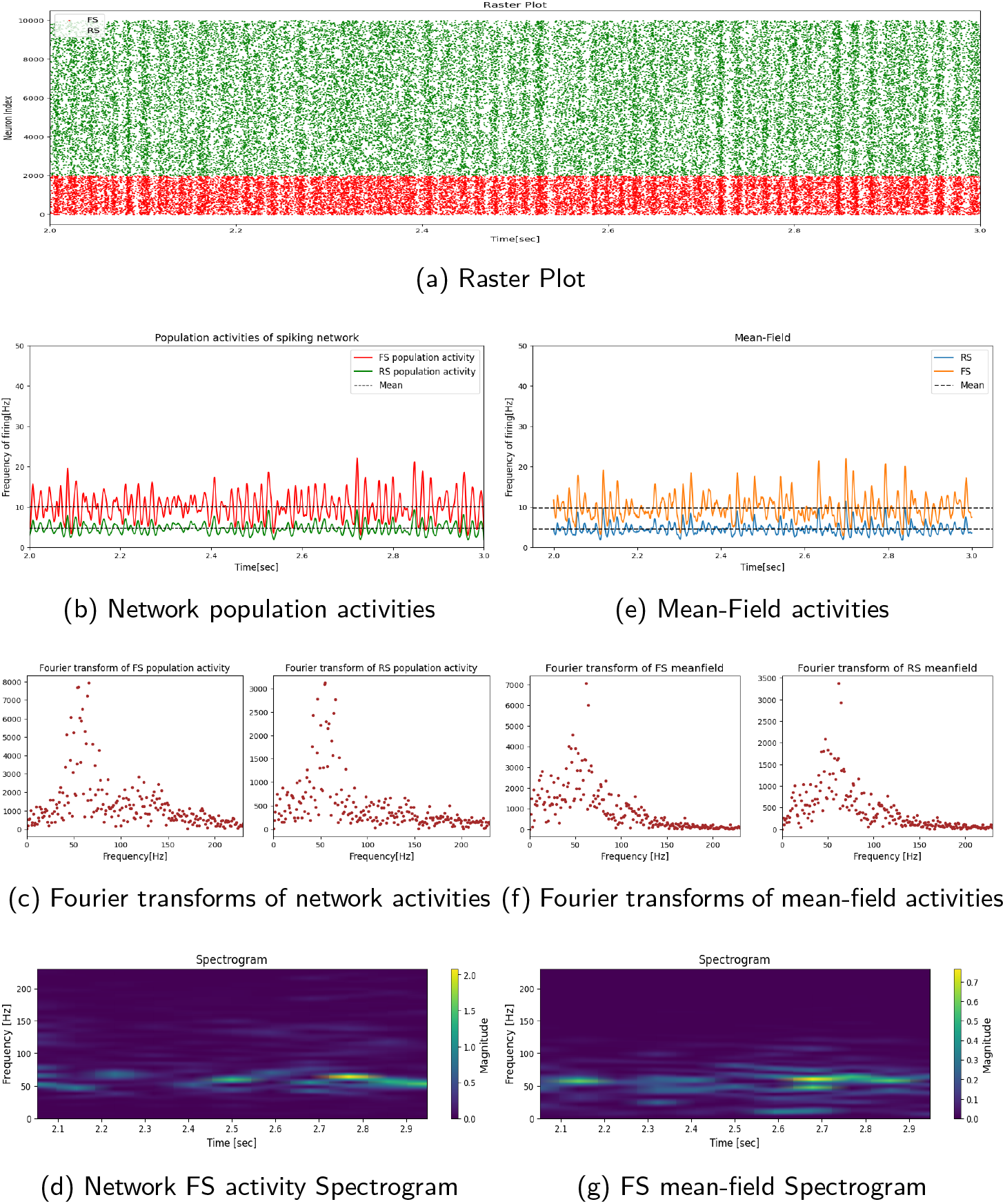
Type 2 spiking network and its corresponding mean-field with 2 ms delay. The external inputs on excitatory and inhibitory populations are 4 Hz and 2 Hz, respectively. Same arrangement of panels as for Fig. 1. The system has a prominent oscillation of around 60 Hz which is nearly the same in the spiking network and the mean-field as seen through Spectrograms and Fourier transform curves. Furthermore, the power of oscillations in the network is in agreement with that of mean-field. Additionally, the power of oscillation in the FS population is higher than that of the RS population both in the network and mean-field. Overall, The mean-field model is in good agreement with the spiking network in terms of both population mean values and the oscillation characteristics.

**Figure 7.**
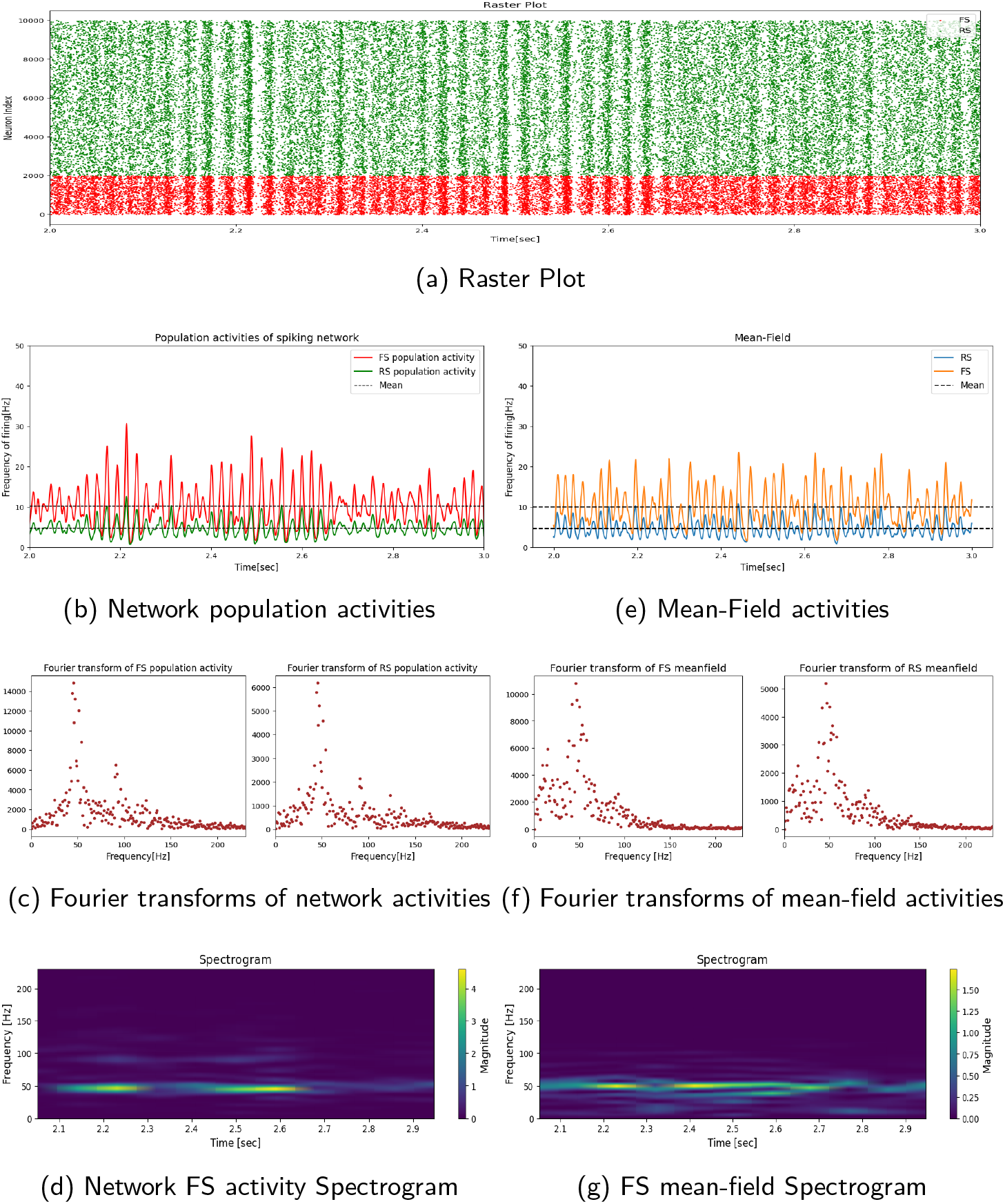
Type 2 spiking network and its corresponding mean-field with 3 ms delay. The external inputs on excitatory and inhibitory populations are 4 Hz and 2 Hz, respectively. Same arrangement of panels as for Fig. 1. The system has a prominent oscillation of around 50 Hz which is nearly the same in the spiking network and the mean-field as seen through Spectrograms and Fourier transform curves. Furthermore, the power of oscillations in the network is in agreement with that of mean-field. Additionally, the power of oscillation in the FS population is higher than that of the RS population both in the network and mean-field. Overall, The mean-field model is in good agreement with the spiking network regarding both population mean values and the oscillation characteristics.

**Figure 8.**
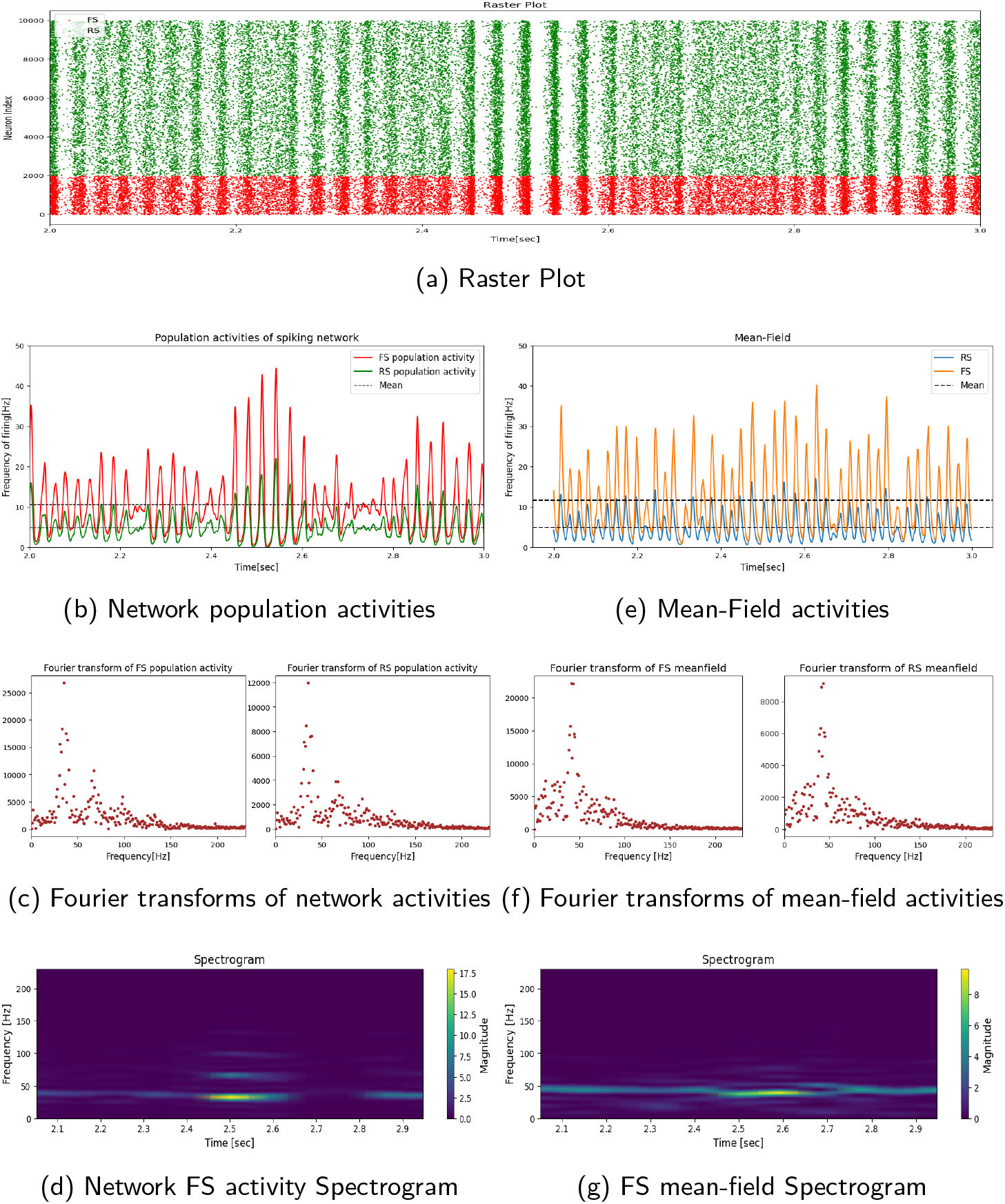
Type 2 spiking network and its corresponding mean-field with 4 ms delay. The external inputs on excitatory and inhibitory populations are 4 Hz and 2 Hz, respectively. Same arrangement of panels as for Fig. 1. The system has a prominent oscillation of around 40 Hz which is nearly the same in the spiking network and the mean-field as seen through Spectrograms and Fourier transform curves. Furthermore, the power of oscillations in the network is in agreement with that of mean-field. Additionally, the power of oscillation in the FS population is higher than that of the RS population both in the network and mean-field. Overall, The mean-field model is in good agreement with the spiking network in terms of both population mean values and the oscillation characteristics.

**Figure 9.**
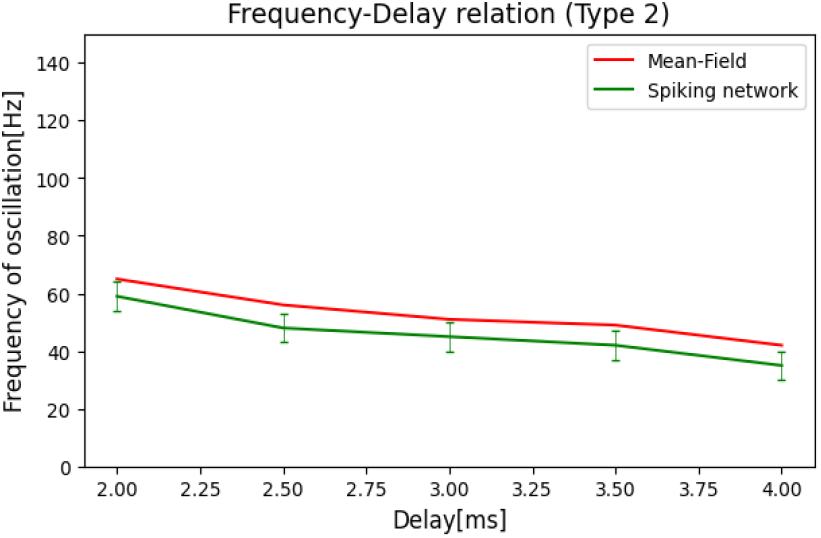
Relation between the frequency of oscillation and the delay for Type 2 networks. The spiking network oscillation frequency matches well with that of the mean-field.

## Discussion

In this study, we have explored spiking networks and respective mean-field models for the generation of gamma oscillations during the AI state in the Cortex and illustrated that they match well with each other. We have studied two types of configurations. In Type 1 networks, the oscillation is generated by the interplay of excitation and inhibition (RS and FS cells in our model), and the synaptic time delay is essential in this mechanism. We have shown that the network oscillation frequency is a decreasing function of the time delay (Fig. 4). It must be noted that here, the synaptic time delay accounts for the axonal and synaptic delays, which may be constant in a local network because the axonal connections are short-range. In the case of large-distance connections, the axonal delay will be proportional to distance, and one could expect more complex behavior in such multiple-time delay systems.

We also showed that for Type 1 networks, the mean-field model, if equipped with time delays, can produce gamma oscillations. The agreement between the oscillation frequency predicted by the mean-field and the spiking network is remarkable (Fig. 4). It must be noted that this agreement is not in contradiction with the Markovian hypothesis which is central in the formalism (El Boustani and Destexhe, 2009). The condition is that the time constant *T* of the meanfield must be smaller, or at maximum comparable to, the oscillation period of gamma oscillations. This may be challenging for the small values of delay, where the oscillation frequency is high (Fig. 4), but astonishingly the mean-field still predicts well the oscillation frequency.

A second type of mechanism was considered for the genesis of gamma oscillations, based on a break-down of the balance between excitation and inhibition. In these Type 2 networks, the external drive was reduced on FS cells. As a result, FS cells exerted less inhibitory overall activity and this produces a different type of gamma oscillation with a lower oscillation frequency (compare Fig. 4 and Fig. 9). Here again, there was a remarkable match between the network frequency and the frequency predicted by the mean-field (Fig. 9). It should be emphasized that the reduction in oscillation frequency in the Type 2 system is due to the increase in overall excitation versus inhibition which also was reported by Brunel and Wang, 2003.

Another interesting feature of Type 2 networks is that they may be relevant to the kind of gamma oscillation induced by drugs like Ketamine. Ketamine is recognized for its ability to trigger significant alterations in brain oscillatory dynamics, which seem to be associated with its antidepressant effects and sensory dissociation (Tian et al., 2023). The electrophysiological characteristics of sub-anesthetic doses of Ketamine typically involve heightened gamma oscillation power alongside reduced delta, alpha, and beta oscillation power (Tian et al., 2023). Even though Ketamine is an antagonist of NMDA receptors, several studies have shown that low doses of Ketamine lead to excitation instead of inhibition (Moghaddam et al. 1997, Su et al. 2018 and Shaw et al. 2015). A previous computational model (Susin and Destexhe, 2023) has modeled this effect by explicitly including NMDA receptors in the RS-FS network. In the present model, we did not have NMDA receptors explicitly, but the decrease of the drive onto FS cells mimics a similar effect of reduced excitation of FS cells and a subsequent dis-inhibition of the network. Thus, the gamma oscillations generated in Type 2 networks are probably of a similar type as Ketamine-induced gamma. Indeed it was shown that Ketamine tends to produce low-frequency gamma oscillations (Shaw et al 2015), very similar to our Type 2 model.

Another aspect of gamma-frequency oscillations is that they tend to occur within an asynchronous-irregular (AI) activity state. This was characterized in humans using micro-electrode recordings (Le Van Quyen et al., 2016). The gamma oscillation appears within the AI state and is more present in FS neurons compared to RS neurons, very similar to our model (compare Fourier plots of RS and FS populations). This study also found that the oscillations in RS and FS populations are globally in phase, as in our model, however, there was a slight phase advance of FS cells compared to RS cells, which our model also exhibits well (see Fig. 10).

**Figure 10.**
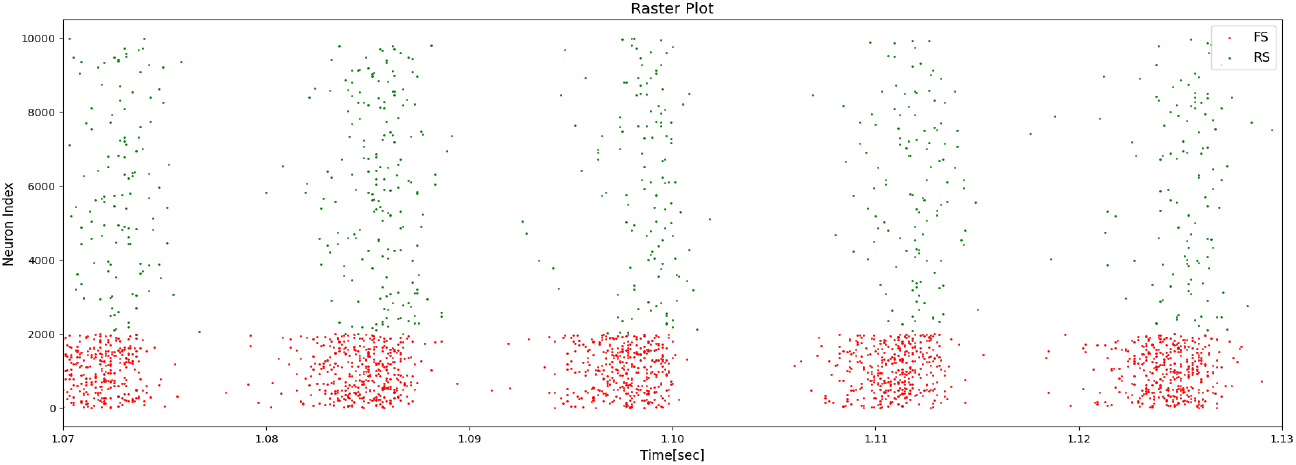
Raster plot of Type 1 network with a synaptic delay of 3 *ms* and an external drive of 5 Hz. Here, all of the synaptic time delays, *exc → exc, exc → inh, inh → exc*, and *inh → inh* are the same without asymmetry. As one can see, there is an obvious phase advance of FS compared to RS cells.

Future studies should investigate several aspects, in continuation of the present work. The main extension is that this model should be ported to larger scales, by integrating gamma-producing mean-fields in large-scale networks of mean-field models (as in Zerlaut et al. 2018). Such an approach can be used to investigate how gamma oscillations organize at large scales in the brain, taking into account long-range connectivity. Since the AdEx mean-field was recently implemented in whole-brain simulations (Goldman et al., 2023), the present mean-field model could form the basis of whole-brain models including gamma-frequency oscillations.

## A Appendix A: Delay asymmetries

In this Appendix, we give more details on the asymmetry in delays that is needed in the mean-field model.

Figure 10 shows a magnified image of a Type 1 network’s raster plot over a gamma cycle. In a gamma cycle, the sequence of spiking activity begins with FS neurons followed by RS cells as can be seen in the figure. This sequential pattern results in a slight phase advance for FS neurons compared to RS cells. Notably, this phenomenon has been experimentally observed in human and primates in vivo (Le Van Quyen et al., 2016).

The mean-field with a constant delay does not robustly generate gamma oscillations, and we needed to consider a slight asymetry of the delays between populations, as *exc → exc* and *inh → inh* equal to (1 + *ϵ*)*τ*_*d*_ while the delays for the other two connections (*exc→inh* and *inh→exc*) equal (1*−ϵ*)*τ*_*d*_. This means that the excitatory recurrent input is received by the excitatory population a little later (2*ϵτ*_*d*_) than the inhibitory input and the excitatory input is received by the inhibitory population a little sooner (2*ϵτ*_*d*_) than the recurrent inhibitory input. This slight asymmetry in delays is compatible with the phenomenon that the FS cells start to spike sooner than the RS cells.

For the spiking network, however, the presence of this asymetry had little impact. The behavior of the spiking network with this asymmetry in delays, is almost similar to the one without the asymmetry. Fig. 11 shows the same network as Fig. 10, but with a delay asymmetry.

**Figure 11.**
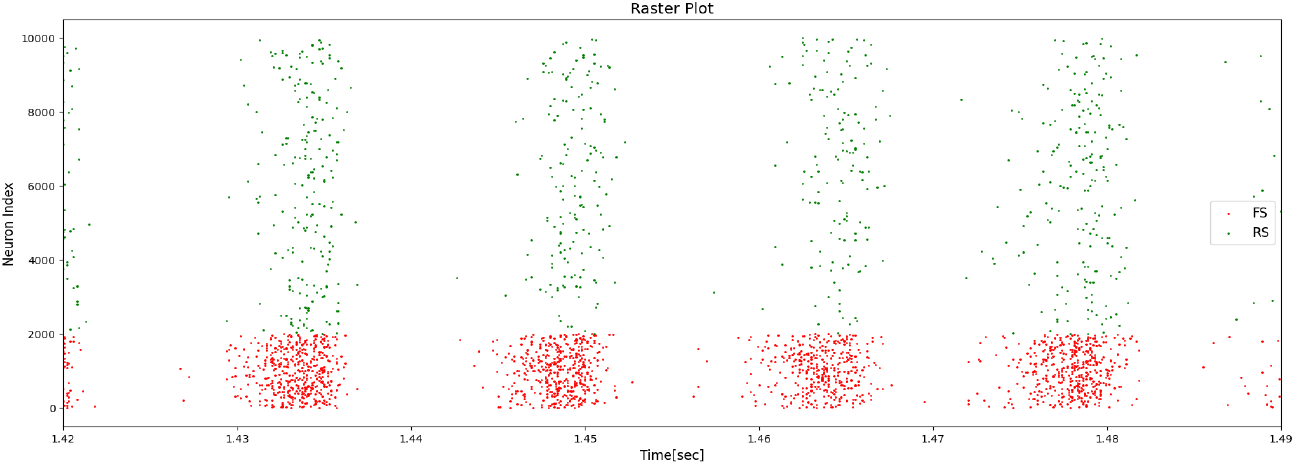
Raster plot similar as in Fig. 10 but with delay asymmetries. *exc → exc* and *inh → inh* delays are (1 + *ϵ*)*τ*_*d*_ while the delays for the other two connections (*exc → inh* and *inh → exc*) equal (1 *− ϵ*)*τ*_*d*_. The phase advance of FS cells compared to RS neurons is similar to the system without delay asymmetries (Fig. 10).

## B Appendix B: Calculation of the transfer function

In this Appendix, we give more details on the calculation of the transfer function in the mean-field model, following a procedure developed by Zerlaut et al. (2016) and di V0lo et al. (2019).

We suppose that the neuron’s firing rate can be expressed as a function of the statistical characteristics of its sub-threshold voltage dynamics. These statistical characteristics are the mean sub-threshold voltage (*μ*_*v*_), its standard deviation (*σ*_*v*_), and the time correlation decay time (*τ*_*v*_).

First we calculate the mean and standard deviation of synaptic conductances which lead to 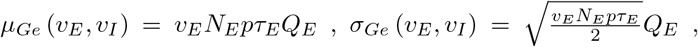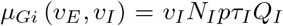, and 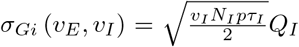 where *N*_*E*_ = 8000 and *N*_*I*_ = 2000 are the number of pre-synaptic excitatory and inhibitory neurons, respectively. Therefore, the input conductance and the effective membrane time constant of a neuron become *μ*_*G*_ (*v*_*E*_, *v*_*I*_ ) = *μ*_*Ge*_ + *μ*_*Gi*_ + *g*_*L*_ and 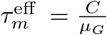. Then assuming that the time scale of the adaptation current is much slower than the time scale of voltage fluctuations and also neglecting the exponential term in Eq.1 we derive the statistics of the membrane sub-threshold voltage as follows.

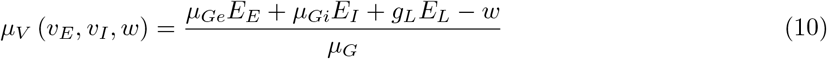

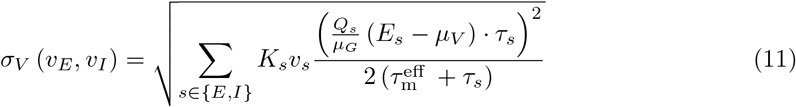

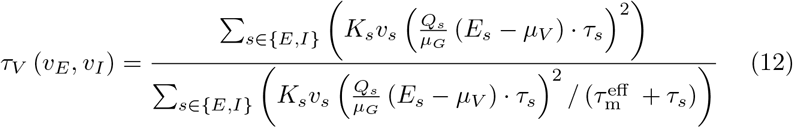

Now one can write the neuron’s output firing rate as

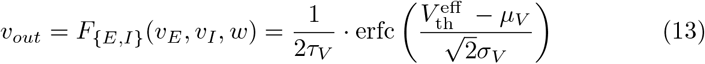

which defines the neuron’s transfer function. Here, erfc is the complementary error function 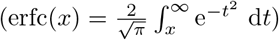 and 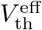 is the effective voltage threshold which can be itself written as a function of *μ*_*V*_, *σ*_*V*_, and *τ*_*V*_ as shown by Zerlaut et al. (2016). Specifically, we take 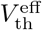 as a second-order polynomial as in Di Volo et al. (2019).

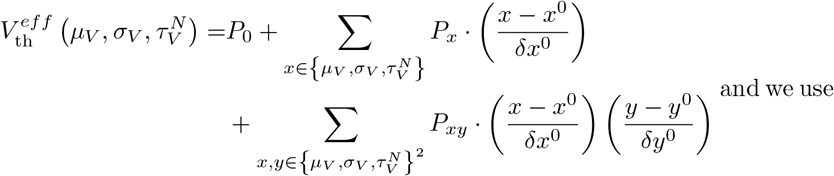

the fitted parameters (Table 3) which were obtained by Di Volo et al., 2019.

**Table 3.**

| Cell Type | $P_0$ | $P_{\mu_V}$ | $P_{\sigma_V}$ | $P_{\tau_V^N}$ | $P_{\mu_V^2}$ | $P_{\sigma_V^2}$ | $P_{(\tau_V^N)^2}$ | $P_{\mu_V \sigma_V}$ | $P_{\mu_V \tau_V^N}$ | $P_{\sigma_V \tau_V^N}$ |
| --- | --- | --- | --- | --- | --- | --- | --- | --- | --- | --- |
| RS-cell | -49.8 | 5.06 | -25 | 1.4 | -0.41 | 10.5 | -36 | 7.4 | 1.2 | -40.7 |
| FS-cell | -51.4 | 4.0 | -8.3 | 0.2 | -0.5 | 1.4 | -14.6 | 4.5 | 2.8 | -15.3 |

## Declarations

### Ethical approval

N/A

### Availability of data and materials

All program codes used in this publication will be made available open-access in Zenodo (Tahvili and Destexhe, 2023).

